# STED super-resolution imaging of membrane packing and dynamics by exchangeable polarity-sensitive dyes

**DOI:** 10.1101/2021.06.05.446432

**Authors:** Pablo Carravilla, Anindita Dasgupta, Gaukhar Zhurgenbayeva, Dmytro I. Danylchuk, Andrey S. Klymchenko, Erdinc Sezgin, Christian Eggeling

**Author notes:** To whom correspondence should be addressed, **Corresponding Authors** Pablo Carravilla: Leibniz Institute of Photonic Technology, Albert-Einstein-Strasse 9, 07745 Jena, Germany., Erdinc Sezgin: SciLifeLab (Gamma-3), Tomtebodavägen 23B, 17165 Solna, Sweden.

## Abstract

Understanding the plasma membrane nano-scale organisation and dynamics in living cells requires microscopy techniques with high spatial and temporal resolution, permitting for long acquisition times, and that allow for the quantification of membrane biophysical properties such as lipid ordering. Among the most popular super-resolution techniques, stimulated emission depletion (STED) microscopy offers one of the highest temporal resolutions, ultimately defined by the scanning speed. However, monitoring live processes using STED microscopy is significantly limited by photobleaching, which recently has been circumvented by exchangeable membrane dyes that only temporarily reside in the membrane. Here, we show that NR4A, a polarity-sensitive exchangeable plasma membrane probe based on Nile Red, permits the super-resolved quantification of membrane biophysical parameters in real time with high temporal and spatial resolution as well as long acquisition times. The potential of this polarity-sensitive exchangeable dye is showcased by live-cell real-time 3D-STED recordings of bleb formation and lipid exchange during membrane fusion, as well as by STED-fluorescence correlation spectroscopy (STED-FCS) experiments for the simultaneous quantification of membrane dynamics and lipid packing, which correlate in model and live-cell membranes.

---

Living organisms meticulously adapt their membrane physical and chemical properties^1^. These changes occur at different temporal and spatial scales; for example, cells can adapt their whole lipidome upon differentiation in a process that takes days^2^; some enveloped viruses like HIV-1 assemble within minutes in sub-micrometric budding sites with a distinct lipid composition^3,4^, and lipid molecules can establish millisecond-lived interactions within nanometric trapping sites^5^. Fluorescence microscopy is a powerful tool to study cell membranes, and by combining it with spectroscopy methods, a number of membrane biophysical parameters can be quantified, such as molecular mobility by fluorescence correlation spectroscopy (FCS)^6^, lipid packing using polarity-sensitive dyes in combination with observation of spectral shifts in fluorescence emission^7^ or membrane tension by measuring fluorescence lifetime^8^. Unfortunately, no fluorescence microscopy technique could so far access the whole range of temporal and spatial scales, i.e. combining super-resolution with long acquisition times and simultaneous readout of biophysical parameters, and new versatile approaches together with improved and sensitive fluorescent probes are required to study biological membranes.

Light diffraction limits the spatial resolution of optical microscopes. This was circumvented by the super-resolution microscopy (SRM) techniques, which have been extensively applied to membrane research^9^. However, SRM introduced new limitations to temporal resolution; for instance, single-molecule localisation microscopy (SMLM)^10^ and MINFLUX imaging^11^ rely on individual molecules emitting photons at different times, and thus require prolonged acquisitions for high localisation precision. Stimulated-emission depletion (STED) microscopy^12^ is, in principle, compatible with time-lapse acquisitions allowing STED microscopy to be combined with spectroscopic techniques, such as (STED-)FCS^5^, raster image correlation spectroscopy^13^, or spectrally-resolved imaging ^14,15^. Unfortunately, although the STED phenomenon is reversible and in theory does not induce enhanced dye destruction (but rather a reduction)^16^, photobleaching rates are above those of conventional microscopy, especially at high STED laser powers required to achieve the highest spatial resolution^17,18^. This higher photobleaching rate seems to be associated with transitions to the more reactive higher excited singlet and triplet states^19,20^ through absorption of STED photons^17,18,21^. Photobleaching remains one of the major complications of STED microscopy, despite big efforts to avoid it, such as the development of novel photostable dyes^22,23^, separation of excitation pulses^24^, the use of high scanning rates^25,26^, or adaptive illumination modes^27^.

An alternative method to circumvent photobleaching is the use of exchangeable dyes. Exchangeable dyes transiently bind to their target and are subsequently removed, thus minimising the photobleaching probability and eliminating photobleached molecules. Exchangeable dyes are the basis for PAINT (point accumulation for imaging in nanoscale topography)^28^, which has been applied for spectral and temporal multiplexing in SMLM^29^. Interestingly, the PAINT concept was introduced by exploiting the transient binding of the exchangeable Nile Red dye to lipid membranes^28,30^. Since then, Nile Red has been used in combination with spectrally resolved SMLM for quantitative cell membrane packing imaging^31–34^. In a recent work, Spahn *et al* demonstrated the potential use of exchangeable dyes in STED microscopy by imaging live bacterial and eukaryotic membranes with Nile Red over the course of minutes^35^. The efficacy of Nile Red as an exchangeable dye relies on its polarity-sensitive nature^36^: because its quantum yield in aqueous solutions is much lower than in hydrophobic environments^37^, only those Nile Red molecules bound to membranes are effectively visualised^28^.

Polarity-sensitive dyes, also known as solvatochromic dyes, change their emission spectra in response to the polarity of the environment due to the dipolar relaxation effect^38,39^. In a membrane context, polarity is directly related to the presence of water molecules at the bilayer interface, which is determined by lipid packing^40^ (Figure 1A). Thus, polarity-sensitive dyes have been extensively used to study the molecular order of membranes and their phase behaviour, i.e. their organization into environments of high (liquid-ordered, Lo) and low (liquid-disordered, Ld) lipid packing^41^. Laurdan was the first polarity-sensitive dye widely used to study cell membranes, due to its high sensitivity to packing changes^42^ and its homogeneous distribution within ordered and disordered environments. However, high susceptibility to photobleaching, Laurdan’s incompatibility with SRM, and an extremely high internalisation rate which hinders plasma membrane staining (a limitation shared with Nile Red), have motivated researchers to find alternative dyes to investigate plasma membranes, such as di-4-ANEPPDHQ^43,44^ or new derivatives of Laurdan^45^ and Nile Red^34,46^.

**Figure 1.**
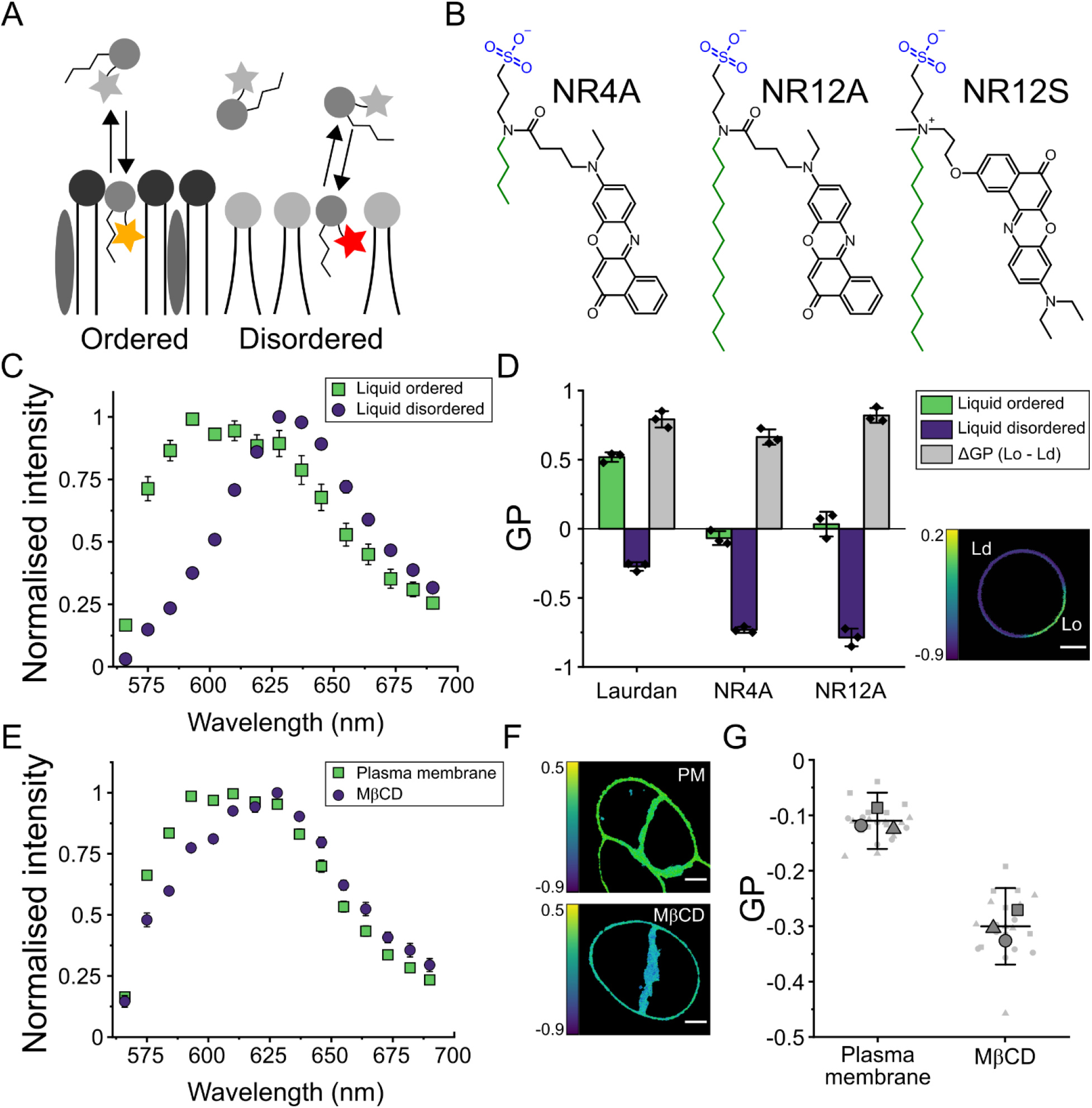
The exchangeable dye NR4A can sense lipid packing changes in model and cell plasma membranes. (A) Exchangeable dyes transiently partition to biological membranes, where their fluorescence spectrum is dependent on lipid packing. (B) Dyes used in this study. Sulfonate groups are marked in blue and alkyl chains in green. Due to its short alkyl chain, NR4A is an exchangeable dye, while NR12A irreversibly binds to membranes. NR4A and NR12A were designed based on NR12S. (C) The emission spectrum of NR4A shifts in response to lipid packing. Phase-separated GUVs were labelled with 250 nM NR4A and imaged by confocal microscopy using spectral detectors. (D) GP analysis of spectral imaging of phase-separated GUVs stained with Laurdan (250 nM), NR4A (250 nM) and NR12A (20 nM). ΔGP is the difference between GP values measured in Lo and Ld environments. The points represent the mean GP of at least 5 GUVs in each of three independent experiments. The micrograph shows the GP image of a phase-separated GUV stained with NR4A. (E) Spectral imaging of living C2BBe1 cells shows that the NR4A emission spectrum undergoes a red-shift upon cholesterol depletion induced by 20 mM MβCD treatment for 1h. (F) GP images of control (PM, top) and MβCD-treated plasma membranes (bottom). GP values were calculated from spectral imaging data. (G) Quantification of plasma membrane GP values shows a significant GP difference upon MβCD treatment. Big symbols are the mean GP value of at least 5 cells for each independent experiment, whereas small symbols represent the GP value of each image. Each symbol (square, triangle, and circle) represents an independent experiment. All measurements were performed at room temperature. In all panels, whiskers are the standard deviation of three independent experiments and scale bars are 5 μm.

To investigate spectral changes in combination with microscopy, in a typical experiment, emitted photons are spectrally separated and collected by two detectors, the lower (blue-shifted) and higher (red-shifted) wavelength signals corresponding to the ordered and disordered channels, respectively. The generalised polarisation (GP) ratiometric function (see Methods section)^40^ is then used to quantify lipid packing^7,47^, with higher GP values corresponding to ordered membranes, and lower values to loosely packed lipid environments. In an alternative approach, spectral detectors available in some commercial microscopes can be used to measure the emission spectra of polarity-sensitive dyes in every pixel, combining the advantages of microscopy and spectroscopy^48^.

Here, we characterise the spectral properties and STED microscopy performance of NR4A, a recently developed Nile Red exchangeable derivative^34^, which selectively stains plasma membranes. We found that because of its enhanced brightness, NR4A shows superior resolution and performance in STED and STED-FCS measurements. Due to its exchangeable nature, NR4A circumvents photobleaching-induced signal loss in STED microscopy experiments and permits long-term super-resolved quantification of plasma membrane biophysical properties, namely lipid packing and dynamics. We showcase possible applications of NR4A by using 3D-STED to image giant plasma membrane vesicle (GPMV) formation and lipid exchange during membrane fusion.

## RESULTS AND DISCUSSION

### NR4A spectral imaging can resolve distinct lipid compositions in model and cell membranes

NR4A was recently developed by linking Nile Red to a plasma membrane targeting moiety which included an anionic sulfonate head (Figure 1B, blue) and a four-carbon alkyl chain (Figure 1B, green). The sulfonate group decreased cell permeability by decreasing the flip-flop rate through the membrane^34,45^, whereas the short alkyl chain permitted transient binding to lipid bilayers^34^. NR12A, a Nile Red-based analogue of NR4A but with a 12-carbon alkyl chain (Figure 1B), showed no exchangeable behaviour^34^ and was used as a control in our experiments. NR4A and NR12A were designed based on NR12S (Figure 1B), a Nile Red-based plasma membrane dye containing a phenolic oxygen, which was hypothesised to cause reduced photostability^34^.

To investigate the sensitivity of NR4A to lipid packing changes, we prepared phase-separated giant unilamellar vesicles (GUVs) made of phosphatidylcholine (DOPC), sphingomyelin and cholesterol (2:2:1 mol ratio), and stained them with NR4A. Confocal spectral imaging revealed a clear red-shift of the emission spectrum in Ld environments as compared to sphingomyelin- and cholesterol-rich Lo environments, with the emission maximum shifting by ca. 35 nm (Figure 1C), as it did for NR12A (Supplementary Figure 1A). To assess whether NR4A’s exchangeable nature influenced its sensitivity to lipid packing, we then quantified the lipid packing resolution as measured by the GP difference (ΔGP) between Lo and Ld environments in phase-separated GUVs (Figure 1D) and compared it with Laurdan, a good standard for sensitivity. NR4A showed a ΔGP (0.66 ± 0.06) comparable to that of NR12A (0.82 ± 0.05) or Laurdan (0.79 ± 0.06). These results were independent of the excitation wavelength (Supplementary Figure 1C-E).

Spectral imaging experiments showed that NR4A is, besides hardly internalizing into the cell (as further outlined later), sensitive to lipid changes in the plasma membrane of living cells too, exemplified by cholesterol depletion by methyl-β-cyclodextrin (MβCD)-treatment of living epithelial C2BBe1 cells (Figure 1E). A comparable red shift was measured for NR12A (Supplementary Figure 1B). Using spectral imaging data for calculation of GP images^48^, where the GP value for each pixel is represented in a colour-coded scale (Figure 1F), we also observed a significant change in plasma membrane packing (Figure 1G).

These results demonstrate that NR4A, in line with the earlier report^34^, is a polarity-sensitive membrane dye that can resolve distinct lipid packing environments in model and cell membranes, with a performance comparable to that of well-established polarity-sensitive dyes, such as Laurdan.

### NR4A enables prolonged time-lapse STED acquisitions of live-cell plasma membranes

To assess whether the exchangeable nature of NR4A could circumvent photobleaching-induced signal loss in STED microscopy measurements, we stained live Ptk2 epithelial cell plasma membranes with NR4A and its non-exchangeable counterpart NR12A. Quantification of the mean intensity recorded upon STED imaging showed a constant signal for NR4A, whereas NR12A was readily photobleached (Figure 2A-B, Supplementary Movies 1-2). Note that the NR12A signal does not reach zero levels, because during the frame-time (10 s) new NR12A molecules are allowed to diffuse in the field of view. The NR4A intensity was constant for longer than 15 minutes, however due to phototoxic effects, signs of cell death could be observed short after 10 minutes.

**Figure 2.**
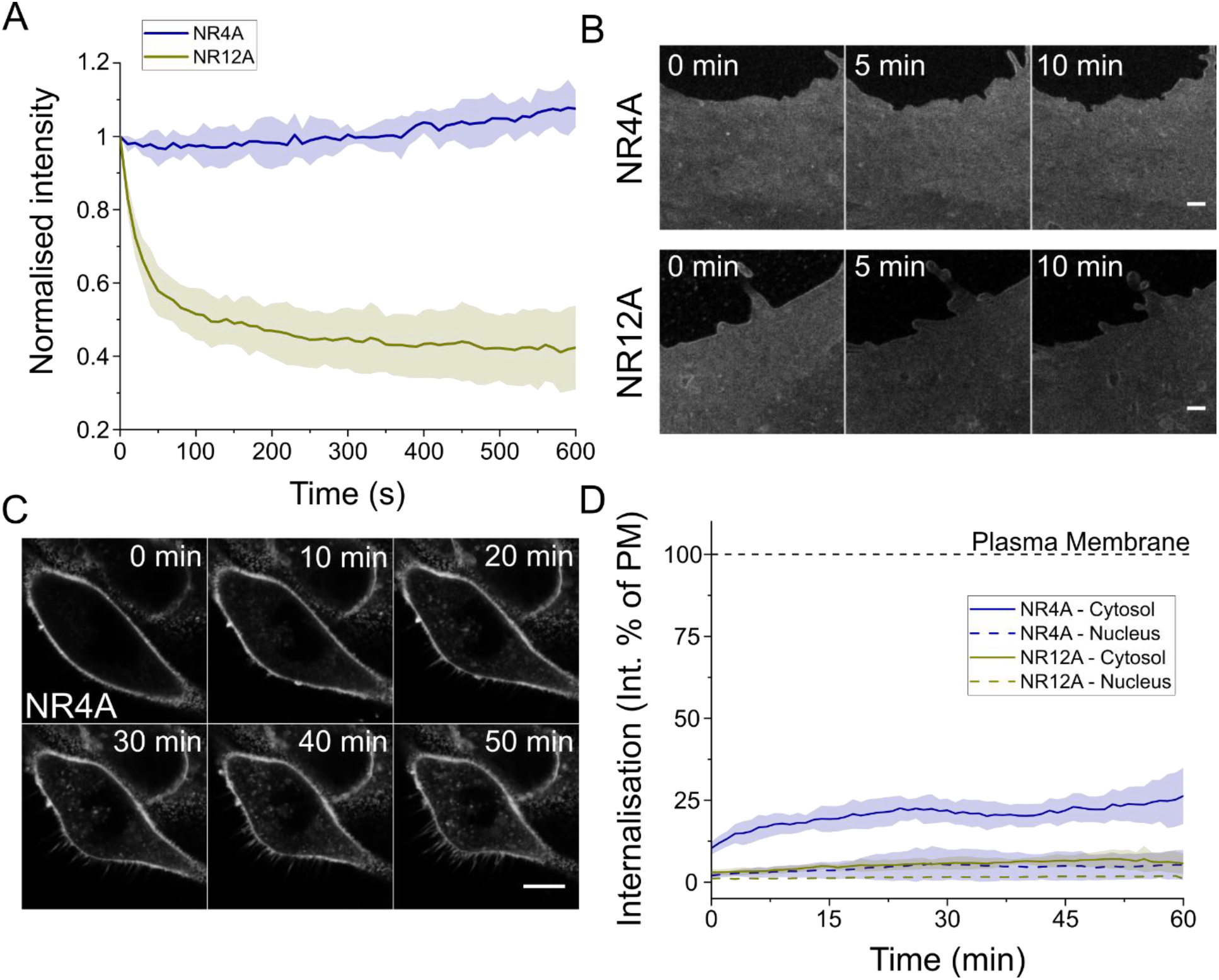
STED imaging of live cell plasma membranes without signal loss. (A) NR4A shows a constant intensity signal during super-resolution STED imaging of Ptk2 plasma membranes over the span of minutes. The NR12A signal decreased due to photobleaching induced by the STED laser. The STED laser power was 250 mW at the sample plane. Normalised intensity is the mean intensity in a 2×2 μm^2^ area divided by the initial mean intensity. The coloured area corresponds to the standard deviation of three independent experiments. Three cells per condition were measured in each independent experiment. (B) Micrographs of STED images of Ptk2 cell plasma membranes illustrating the absence of photobleaching of NR4A as compared to NR12A. Scale bar is 1 μm. (C) NR4A selectively stains live cell plasma membranes. Micrographs of C2BBe1 cells upon incubation with NR4A acquired in confocal mode. (D) Quantification of NR4A and NR12A internalisation. Internalisation is measured as the intensity in regions of interest which could unequivocally be identified as the cytosol or the nucleus divided by the intensity signal recorded at the plasma membrane. The coloured area corresponds to the standard deviation of three independent experiments. Dashed line marks 100%, which corresponds to the plasma membrane signal. All measurements were performed at room temperature.

Previously, 90-100 nm resolution was obtained using NR12S with STED imaging^14^. In line with the reported increased brightness and photostability of NR4A and NR12A as compared to NR12S^34^, NR12A showed a significantly higher raw intensity in STED time-lapse acquisitions of Ptk2 cell membranes, under the same imaging conditions and concentration compared to NR12S (Supplementary Figure 2A). Thus, even if NR12A is readily photobleached in STED experiments, it still shows superior performance compared to NR12S. Importantly, lipid packing quantification, showed that the measured GP increased upon NR12A photobleaching, while it remained constant for NR4A (Supplementary Figure 2B). This phenomenon can presumably be caused by photobleached dyes reacting with surrounding lipids^42,49^, which might be prevented in the case of NR4A due to transient binding and thus lower reaction probability.

Another important limitation when studying plasma membranes is dye internalisation. This problem is aggravated when imaging polarity-sensitive dyes, since the molecular order of inner organelle membranes is significantly lower than that of the plasma membrane^50^, and thus dye internalisation induces a change in the emission spectrum. This is the case of Nile Red, which can be exploited to image inner organelle membranes^35^, but might report an underestimated packing value when studying plasma membranes. NR4A, NR12A and NR12S were designed to prevent dye internalisation by including anionic sulfonate or zwitterionic groups^34,46^. Sulfonate (NR4A, NR12A) and zwitterionic (NR12S) groups reduce transmembrane translocations (flip-flop) of membrane dyes, keeping them in the outer leaflet of plasma membranes^45,46^. To measure NR4A internalisation, we stained C2BBe1 human epithelial cells and quantified the intensity recorded in three distinct cell regions, namely the plasma membrane, the cytosol, and the nucleus. Even after 50 minutes, most of the NR4A signal was detected at plasma membranes (Figure 2C-D, Supplementary Movie 3). However, the cytosolic signal of NR4A was significantly higher than that of NR12A (Figure 2D), which almost exclusively remained in the plasma membrane for the whole duration of the experiment (Supplementary Figure 2C, Supplementary Movie 4). We hypothesize that the highly concentrated unbound pool of NR4A in the extracellular medium could be encapsulated and transported through endocytosis inside the cell, where it would partition to inner membranes.

Our results show that NR4A can be used to perform time-lapse imaging and quantification of lipid packing of live cells using super-resolution STED microscopy, circumventing photobleaching-induced signal loss. In combination with selective plasma membrane staining, this makes NR4A an appropriate dye for time-lapse STED studies. We argue that although NR12A suffers from photobleaching, it can still be used for experiments where continuous illumination or high STED powers are not required. Among the advantages of NR12A are its higher selectivity for plasma membranes (Figure 2D), and the need to use significantly lower concentrations (10-20 nM) as compared to NR4A (250-500 nM), due to NR12A’s strict partitioning to membranes^34^.

### STED-FCS measurements show superior resolution of NR4A and NR12A compared to NR12S

STED microscopy resolution is determined by the photophysics of the imaged fluorophore, and thus it has to be measured for each dye independently. Accurately determining the resolution of STED microscopy can be challenging, especially when imaging continuous structures like membranes, where the distance between two dyes continuously changes. STED-FCS constitutes an alternative approach to accurately quantify resolution^51^. In an FCS experiment, the average time a molecule takes to cross the confocal volume is measured, thus permitting the calculation of its diffusion coefficient. By measuring the transition time at different STED laser powers on a supported lipid bilayer (SLB), we can calculate the volume of the observation spot for each STED power^51^. Since an SLB is a 2-dimensional structure, this approach reports on the size of the observation spot in the *xy* dimensions (ω_0_, see Methods section for more details).

To quantify the resolution offered by NR4A, we performed STED-FCS measurements of DOPC SLBs. NR4A and NR12A showed a maximal resolution of 80.4 ± 1.4 nm and 77.1 ± 1.4 nm respectively, at 250 mW STED laser power (Figure 3A). This resolution is significantly higher than that obtained for NR12S under the same imaging conditions (95.6 ± 7.4 nm). The major difference between NR4A/NR12A, and the original NR12S, is the reverse orientation of Nile Red in the bilayer and the lack of a phenolic oxygen in the fluorophore of NR4A/NR12 (Figure 1B), which was hypothesised to cause lower photostability^34^. To quantify the effect of this modification in molecular brightness, we measured the counts per molecule of the three dyes at different STED powers using STED-FCS (Figure 3B). As expected, NR4A and NR12A showed a comparable brightness, approximately 3 times higher than that of NR12S, which could account for the observed resolution difference. Moreover, STED-FCS curves obtained for NR4A/NR12A showed significantly less noise compared to those measured with NR12S (Figure 3C), as expected for brighter molecules.

**Figure 3.**
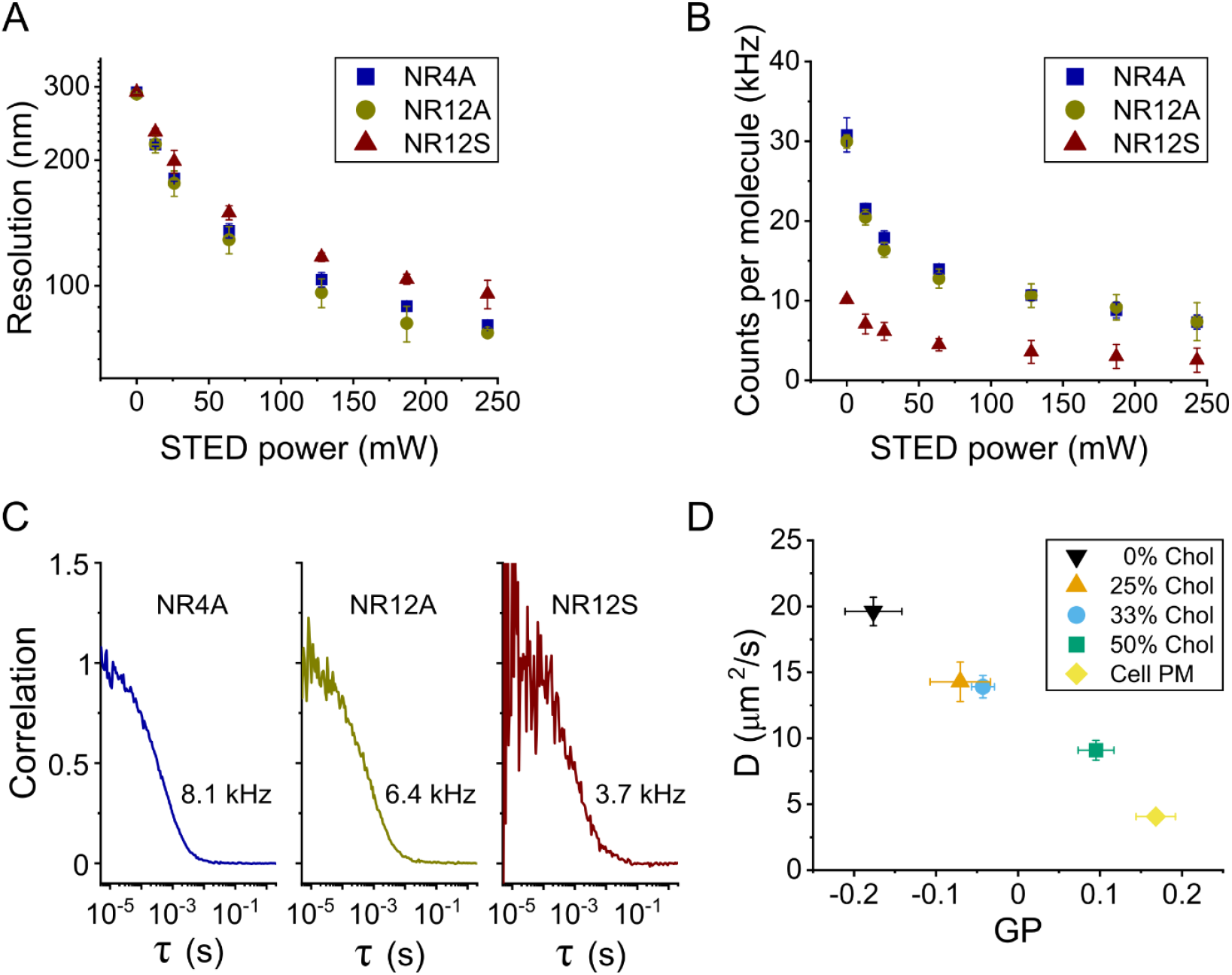
STED-FCS measurements of membrane dynamics and packing. (A) NR4A and NR12A offer higher resolution that NR12S as quantified by STED-FCS measurements on DOPC SLBs. (B) NR4A and NR12A also show superior brightness at all measured resolutions. (C) Exemplary STED-FCS curves obtained on DOPC:cholesterol (1:1 mol ratio) SLBs at 250 mW STED laser power. The average counts per molecule of those particular measurements are indicated. (D) Simultaneous quantification of the diffusion coefficient (D) of NR4A, and its surrounding lipid packing show a negative correlation between packing and dynamics, as measured at 25 mW STED power in cholesterol containing SLBs and in the plasma membrane of live Ptk2 cells. The observed correlation is maintained independent of the STED power (Supplementary Figure 3B). All measurements were performed at room temperature. In all panels, whiskers are the standard deviation of at least three independent measurements (at least 5 acquisitions per condition per independent measurement).

In an attempt to simultaneously measure membrane dynamics and packing, we combined STED-FCS acquisitions with GP analysis. For that, we prepared SLBs containing increasing amounts of cholesterol and measured the intensity fluctuations of NR4A in two different spectral channels (580-625 nm and 650-700 nm) at different resolutions (i.e. STED laser powers). We measured the diffusion coefficient of NR4A using FCS, and quantified lipid packing by calculating the GP function from the average photon count measured in each channel. We observed a linear correlation between lipid packing and diffusion at different cholesterol concentrations (Figure 3D), which was independent of the resolution (Supplementary Figure 3B). The observed correlation was maintained even when including data acquired on the plasma membrane of live Ptk2 epithelial cells (Figure 3D). It must be noted that GP values strongly depend on the STED laser power^14^ (Supplementary Figure 3A), although the GP-resolution (dynamic range) was maintained at all measured STED powers. Thus, STED-GP values can only be compared to those obtained under the same imaging conditions (e.g, same STED power and dye concentration).

Altogether, these results show that, due to their increased brightness, NR4A and NR12A offer superior resolution and performance when compared to the previous generation counterpart NR12S in STED, FCS and STED-FCS experiments. Despite only transiently partitioning to membranes, the binding time for NR4A is longer than the time it takes to cross the observation spot (ca. 2 ms in cell plasma membranes), making it compatible with FCS measurements. Two-channel FCS acquisitions permit the simultaneous calculation of diffusion coefficients and lipid packing by means of the GP function. By employing this approach, we found that the lipid packing and bilayer dynamics anticorrelate in model and live-cell membranes, suggesting that the slow diffusion coefficients commonly measured in plasma membranes can be ascribed to the high degree of lipid packing of this structure, which arises from its distinct lipid composition^50^, as previously suggested for highly packed viral membranes^52^.

### Live 3D-STED imaging of cell membranes

We then proceeded to assess whether NR4A could distinguish lipid packing changes in live cells using STED microscopy. Since our current STED setups were not equipped with spectral detectors, we performed classic two-channel acquisitions using emission filters typically found in STED setups (580-630 nm for the ordered channel and 650-700 nm for the disordered channel). We found that this combination of filters offered a high GP-resolution that permitted distinguishing control and cholesterol-depleted (MβCD-treated) plasma membranes at high STED powers (Figure 4A) and in confocal mode (Supplementary Figure 4A).

**Figure 4.**
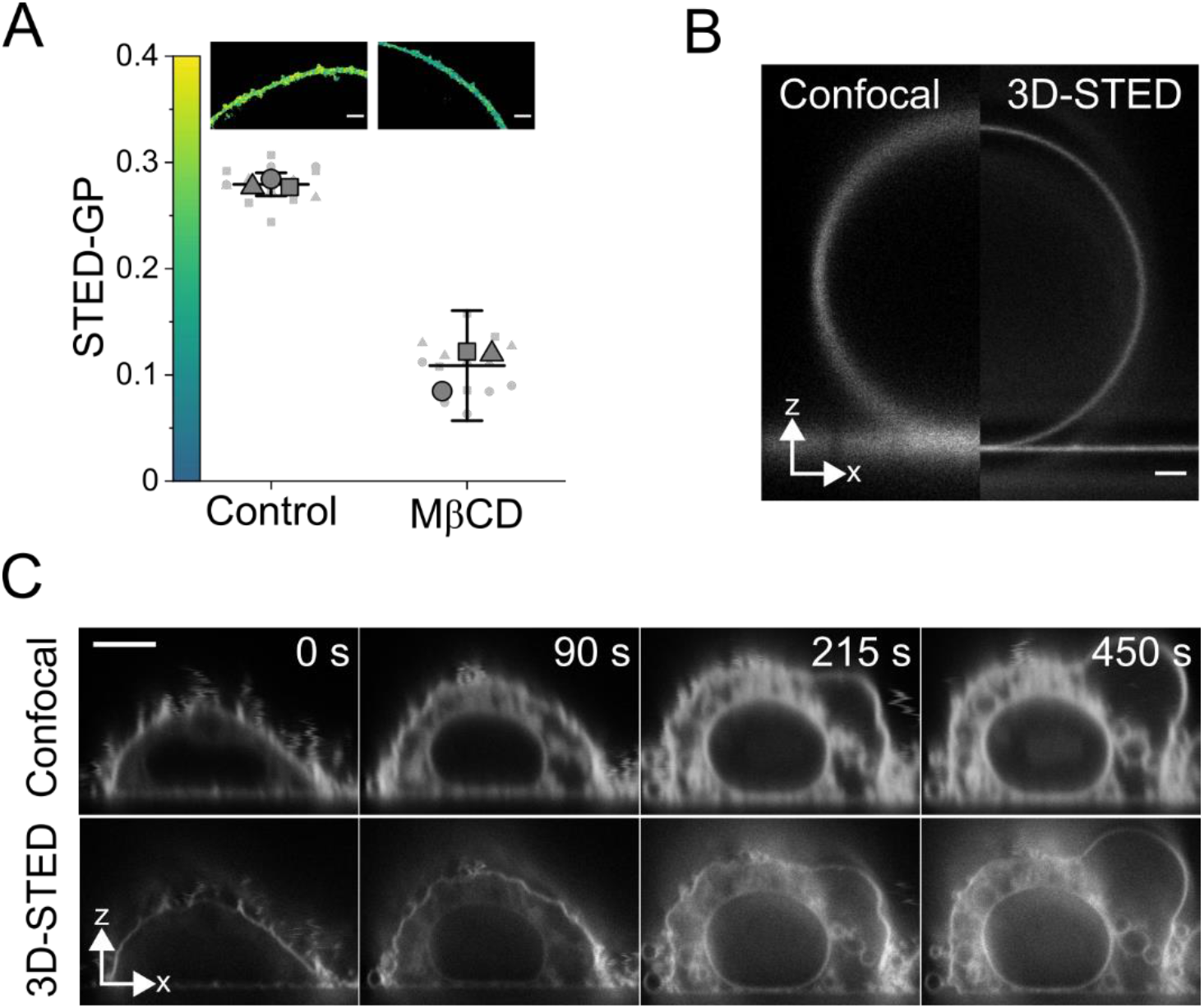
3D-STED visualization of live cell plasma membranes. (A) STED-GP quantification of cholesterol-depletion induced packing changes in C2BBe1 cells. STED laser power was 250 mW. Each big symbol is the mean GP value of each independent experiment (at least 5 cells), whereas small symbols represent the GP value of each image. Symbols (square, triangle, and circle) represent independent experiments. Lines represent the mean and whiskers the standard deviation of three independent experiments. (B) Confocal and 3D-STED microscopy micrograph of a DOPC GUV labelled with 200 nM NR4A. Scale bar is 1 μm. (C) GPMV formation induced by DTT/PFA treatment of CHO cell plasma membranes as visualized by NR4A confocal (top) and 3D-STED (bottom) imaging. Due to DTT/PFA-induced membrane disruption, NR4A can diffuse inside the cell. NR4A concentration was 500 nM. Scale bar is 5 μm. All measurements were performed at room temperature.

In 3D-STED, also known as z-STED, the resolution is increased in three dimensions (*xyz*) by using a “bottle-shaped” depletion scheme instead of the classic “doughnut-shaped” beam^53^. 3D-STED was recently shown to be a powerful tool to investigate living membranes^15^. Here, we wanted to prove the potential of combining 3D-STED and exchangeable dyes for membrane imaging. 3D-STED microscopy of artificial membranes stained with NR4A clearly showed the increase in lateral and axial resolution (Figure 4B). As a model to investigate membrane changes in real time, we induced GPMV formation (i.e. plasma membrane vesiculation) with dithiothreitol (DTT) and paraformaldehyde (PFA)^54^. Plasma membrane vesiculation could be readily observed in CHO cells (Figure 4C, Supplementary Movie 5) and epithelial human 293T cells (Supplementary Figure 4B). The improved axial resolution offered by 3D-STED permitted the visualisation of vesiculation and swelling of inner organelles (Figure 4C). Unfortunately, DTT/PFA treatment induced permeabilization of plasma membranes to NR4A, as demonstrated by the immediate staining of inner membranes (Figure 4C), while untreated cells showed a significantly lower cytosolic signal (Supplementary Figure 4C). GPMVs have been shown to be permeable to small solutes, however cell plasma membranes remain impermeable during GPMV formation^55^. In line with these results, DTT/PFA treatment did not induce plasma membrane permeabilization to hydrophilic solutes, while NR4A got internalised (Supplementary Figure 4D), hampering a detailed plasma membrane packing analysis during GPMV formation.

### Super-resolved live monitoring of lipid packing during membrane fusion

*In vitro* studies of membrane remodelling processes are among the main applications of polarity-sensitive dyes. Real-time monitoring of such phenomena has so far been hampered by photobleaching. Taking advantage of the exchangeable nature of NR4A, we decided to investigate heterotypic membrane fusion, i.e. the merger between two membranes of distinct composition. For that, we prepared SLBs made of DOPC, phosphatidylethanolamine (DOPE) and phosphatidylserine (DOPS) (4:3:3 mol ratio), and incubated them with GUVs made of palmitoyl, oleoyl-phosphatidylcholine (POPC) and cholesterol (2:1 mol ratio). We simultaneously labelled the SLB and GUVs by adding 500 nM NR4A. Both membrane environments could be distinguished by their different lipid packing in confocal measurements, as reported by the GP parameter (Figure 5A, top left panel). Subsequent addition of 10 mM calcium chloride readily promoted membrane fusion (Figure 5A, Supplementary Movie 6), in line with the well-characterised fusogenic activity of divalent cations^56^. By monitoring the process in real-time, we could differentiate the following steps (Supplementary Movie 6): (i) contact, the vesicle is brought in close contact to the supported bilayer, the GUV membrane is highly mobile; (ii) hemifusion, the vesicle is docked, lipid exchange can be observed as reported by the GP change (Figure 5B); and (iii) full fusion, both membranes form a continuous bilayer. Since NR4A only partitions to the outer leaflet of bilayers, complete GP equilibrium could be observed after hemifusion.

**Figure 5.**
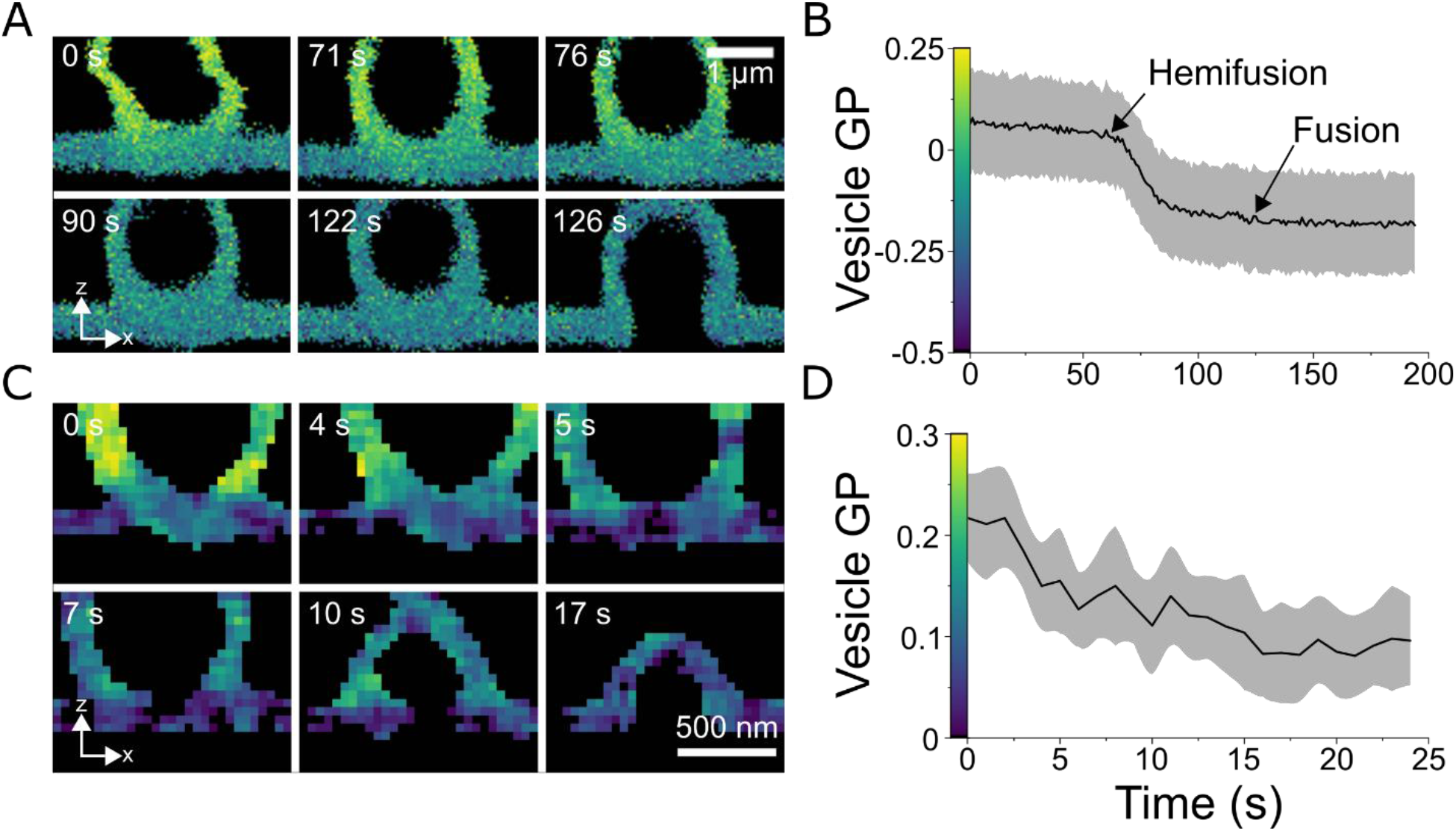
Real-time visualization of membrane fusion by confocal and 3D-STED microscopy of NR4A. (A) *XZ* GP micrographs of a confocal time-lapse of a GUV made of DOPC:DOPE:DOPS (4:3:3 mol ratio) and a POPC:Chol (2:1 mol ratio) SLB. 10 mM calcium addition promoted membrane fusion, that occurred after hemifusion and lipid exchange. GP changes of the GUV over time are shown in B. (C) Membrane fusion and lipid packing monitoring of a sub-micrometre-sized vesicle. Lipid compositions are as in A. (D) Vesicle lipid packing change over time. Scale bars are 1 μm in A and 500 nm in C, as indicated. In B and D, the greyed area is the standard deviation of the vesicle GP in A and C, respectively. All measurements were performed at room temperature.

Endo- and exocytosis involve significantly smaller vesicles than GUVs. Therefore, we next investigated the fusion of submicrometric-sized vesicles with SLBs, which could be resolved by 3D-STED GP imaging (Figure 5C, Supplementary Movie 7). Lipid exchange and fusion happened in a shorter time scale (Figure 5D), due to the smaller size of the vesicles. Finally, we prepared 400 nm large unilamellar vesicles (LUVs), smaller than the axial resolution of confocal microscopy (Supplementary Figure 5A-B, Supplementary Movie 8). In this case, fusion occurred seconds after attachment, and total lipid exchange occurred within two frames (Supplementary Figure 5C-E), which could only be resolved with 3D-STED imaging. In summary, we show that exchangeable polarity-sensitive dyes can be useful tools to image fusion in real time with conventional or super-resolved fluorescence microscopy. Lipid exchange could be quantified by means of GP imaging, resolving the different steps of the fusion process.

### Limitations of this study

Here, NR4A is presented as a STEDable dye that can circumvent photobleaching-induced signal loss. However, STED imaging of plasma and model membranes still presents several obstacles that need to be addressed before attaining the ultimate goal of unlimited live imaging with molecular resolution. First, the resolution offered by NR4A and NR12A is around 80 nm, approximately half of the resolution commonly achieved with live-cell STED microscopy of fluorescent lipid analogues labelled with dyes such as Abberior STAR RED or ATTO647N. Still, to the best of our knowledge, NR4A and NR12A are the polarity-sensitive dyes offering the best STED resolution so far. Moreover, to achieve the highest resolution, STED microscope requires the use of highly focused laser beams with a power in the range of hundreds of mW, which can induce cell photodamage^57^, hampering long-time live-cell studies.

Polarity-sensitive dyes strongly suffer from photoselection, i.e. the preferential excitation of the fluorophore in respect to the excitation light polarisation, which can introduce an underestimation in lipid packing quantification. Since membranes fix the orientation of polarity-sensitive dyes, *xy* imaging needs to be performed with circularly polarised light, while 3D/*z* imaging would require a combination of circularly and axially polarised light, an approach that to the best of our knowledge remains unexplored. Furthermore, Nile Red-based dyes show excitation and emission spectra that overlap with those of the most popular fluorophores and fluorescent proteins, hampering their combined use. The ability to transiently bind to their target structure confers exchangeable dyes their switchable nature, however, since most of the dye remains in solution, the concentrations needed to perform each experiment are at least one order of magnitude higher than that with classic membrane dyes, significantly increasing their cost. Moreover, when used at high concentrations (500 nM), NR4A internalisation can also impose a limitation, if processes that span for longer than an hour are imaged.

## CONCLUSIONS

Here, we studied whether exchangeable probes sensitive to polarity can improve quantitative super-resolution STED imaging of lipid order and dynamics in biomembranes. To this end, we selected NR4A, a Nile Red-based probe previously designed for selective imaging of plasma membranes using SMLM. We found that this exchangeable polarity-sensitive probe can circumvent signal loss induced by dye photobleaching, especially in STED microscopy. It can be used to image model and plasma membranes for prolonged periods of time, while monitoring membrane biophysical properties such as lipid packing or dynamics. NR4A is sensitive to changes on lipid composition in confocal (Figure 1) and STED (Figure 4) microscopy. NR4A permits continuous STED imaging of plasma membranes, which is no longer limited by photobleaching, but rather by cell photodamage (Figure 2). Signal fluctuation analysis methods such as FCS or its super-resolved STED-FCS mode, can simultaneously report on packing and dynamics in model and plasma membranes (Figure 3). NR4A is compatible with 3D-STED, which offers an order of magnitude increase in resolution in all axes, permitting the long-time visualisation of previously unresolvable processes in cells (Figure 4) and model membrane systems (Figure 5). Finally, we would like to highlight the versatility of NR4A, which can be used in confocal, widefield, SMLM^34^, and STED microscopy experiments.

## METHODS

### Dye synthesis

The following fluorescent probes were synthesised according to the previously published procedures: NR4A^34^, NR12A^34^ and NR12S^46^.

### Model membrane systems

DOPC, DOPE, DOPS, POPC and cholesterol were purchased from Avanti Polar Lipids Inc. SLBs were formed by spin-coating^51^. Glass coverslips (#1.5) were immersed in piranha solution (H_2_SO_4_:H_2_O_2_ 3:1 vol ratio) for 2 h, followed by five washing steps with abundant distilled H_2_O. Clean coverslips were stored in H_2_0 for up to a week. Lipid mixtures were prepared in chloroform:methanol (2:1 vol ratio) at a concentration of 1 g/L. 25 μL of the lipid mixture were added to a piranha-cleaned coverslip and immediately (within the same second) spread through spin-coating at 3000 rpm for 30 s. The coverslip was mounted in an Attofluor chamber (Thermo Fisher Scientific) and the resulting lipid film hydrated in HEPES-buffered saline (HBS, 150 mM NaCl, 10 mM HEPES, pH 7.0) and washed thoroughly with HBS 5 times. SLBs were labelled by adding 10-20 nM of NR12A/NR12S or 200 nM NR4A.

GUVs were prepared using the electroformation method. Lipid mixtures were prepared at a final concentration of 1 g/L in chloroform. The lipid mixture (5 μL) was spread on two parallel platinum wires and subsequently dried with a gentle argon stream and 5-minute vacuum. The wires were dipped in a home-made polytetrafluoroethylene (PTFE) chamber filled with 300 mM sucrose. GUVs were formed by connecting the wires to a function generator and applying a 2 V, 10 Hz AC current for 1 h, followed by a 2 V, 2 Hz current for 30 min. GUVs were collected with a 1000 μL plastic pipette tip; to avoid GUV rupture the tip diameter was widened by cutting it approximately 2 cm from the end. GUVs were transferred to a new tube containing 750 μL HBS and left to sink for 15 minutes, due to the higher sucrose density. The bottom 100 μL of the tube were then collected with a cut tip and transferred to a new tube for labelling by adding 10-20 nM NR12A/NR12S. NR4A labelling was attained by adding 200 nM dye directly to the imaging chamber prior to GUV addition. Finally, GUVs were transferred to a bovine serum albumin (BSA)-blocked Ibidi 8-well μ-slide prefilled with 300 μL HBS. The glass surface was blocked prior to GUV addition by incubating the wells with 200 μL 2g/L BSA (Sigma-Aldrich) for 15 minutes, followed by 5 washing steps with 400 μL distilled H_2_O.

LUVs were prepared following the extrusion method. Lipid mixtures in chloroform were dried under an argon stream for 1 minute, followed by vacuum drying for 45 minutes. The resulting lipid films were hydrated in HBS to a final 1 mM concentration, followed by 10 freeze (liquid N_2_, 1 min) and thaw (37 °C water bath, 3 min) cycles and 30 extrusion cycles through two 400 nm filters (Whatman) using a Mini-Extruder (Avanti Polar Lipids Inc).

### Cell lines

C2BBe1 human enterocytes were grown at 37 °C and 5% CO_2_ in Dulbecco’s Modified Eagle’s Medium (DMEM) supplemented with 10% foetal bovine serum (FBS), 1% L-glutamine, 0.01 mg/ml Holo-Transferrin, penicillin, and streptomycin. They were subcultivated at a 1/10 ratio every four days and kept until passage 20. Cells were seeded on collagen I-coated glass coverslips at a final concentration of 1×10^5^ cells/ml in 6-well plates. Cover slides were coated with collagen I at a final concentration of 10 μg/ml, incubated 2 h at room temperature in the dark, and subsequently dried at 37 °C. Ptk2 potoroo kidney cells were grown at 37 °C/5% CO_2_ in DMEM supplemented with 10% FBS, 1% L-glutamine, penicillin, and streptomycin. They were subcultivated at a 1/3 ratio every three days and kept for 30 passages. CHO-K1 Chinese hamster ovary cells were grown at 37 °C and 5% CO_2_ in DMEM-F12 supplemented with 10% FBS. CHO-K1 cells were subcultured at a 1/8 ratio twice per week and kept for 30 passages.

For imaging experiments, cells were grown in 25 mm coverslips for two days. Coverslips were mounted on Attofluor chambers, washed twice with Leibovitz’s L-15 medium (Thermo Fisher Scientific) and afterwards, the corresponding dye was added. In cholesterol depletion experiments, C2BBe1 cells were treated with freshly prepared 20 mM MβCD (or medium in the case of control cells) and incubated for 1 h at 37 °C/5% CO_2_. Finally, cells were washed twice with Leibovitz’s L-15 medium, labelled with 20-50 nM NR12A or 200-500 nM NR4A, and subsequently imaged.

### Spectral Imaging and GP calculation

Spectral images were acquired using a Zeiss 880 confocal laser scanning microscope equipped with a C-APOCHROMAT 40×/1.2 W Korr FCS objective lens (Zeiss). To avoid photoselection, circular polarisation was generated by introducing a phase retarder in the DIC slider of the objective lens. Laurdan (Thermo Fisher Scientific) and NR4A/NR12A were excited using 405 nm and 561 nm laser lines, respectively, with an excitation power of 10 μW at the sample plane. Emitted photons were de-scanned, passed through a 1.0 Airy unit pinhole and collected by a 32-spectral channel GaAsP detector with an 800 V gain. Each spectral channel consisted of an 8.9 nm window. Pixel size was set to 100 nm and the dwell time to 2 us. Images were analysed using Fiji^58^. To calculate emission spectra, an intensity threshold was applied to the images to analyse only pixels corresponding to membranes. In the case of phase-separated GUVs, each phase was analysed separately. For every image, the average intensity signal was then calculated for each channel. To quantify lipid packing the GP value for each pixel was calculated using the GP function^40^:

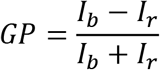

where *I*_*b*_ and *I*_*r*_ are the intensity recorded at 435 nm and 510 nm for Laurdan, and 575 and 640 nm for NR4A/NR12A. GP values were calculated using the GP Plugin^48^ for Fiji. In the case of two-channel images acquired with STED microscopes, an intensity threshold was applied to the sum of both channel images, and GP values for each pixel were calculated using the GP function, where *I*_*b*_ and *I*_*r*_ are the intensities recorded at the ordered (580-630 nm) and disordered (650-700 nm) channels. Mean values for each independent experiment were considered as the mean GP value of at least five images. Graphical representation in Figures 1G and 4A is based on a previous report^59^. For FCS and STED-FCS GP measurements, *I*_*b*_ and *I*_*r*_ are the average photon counts on each channel.

### STED imaging

STED images were acquired in a custom Abberior Expert Line laser scanning STED microscope using an UPlanSApo 60×/1.2 water immersion objective lens equipped with a correction collar (Olympus). Resolution strongly depended on the right adjustment of the collar, especially in 3D-STED mode. NR4A/NR12A were excited by a 561 nm pulsed diode laser PDL-T 561 (Abberior Instruments GmbH) with an excitation power of 10 μW at the sample plane. Fluorescence was inhibited by a 775 nm PFL-40-3000-775-B1R 40 MHz pulsed laser (MPB Communications). The STED beam power at the sample plane for each experiment is specified in figure legends. The beam shape for 2D (doughnut) or 3D depletion (bottle-shape) was created using a spatial light modulator (SLM). To align the STED beam on top of the excitation PSF 4-colour TetraSpeck 100 nm microspheres (Thermo Fisher Scientific) were used as a reference. The STED beam position was corrected in respect to the confocal signal by adjusting the grating of the SLM. The orientation of the depletion beam was fine-tuned by adjusting the SLM offset. Emitted photons were collected through the objective lens, de-scanned, passed through a 1.0 airy unit pinhole, and finally collected by single photon counting SPCM-AQRH-14-TR avalanche photodiodes (Excelitas Technologies) equipped with appropriate filters (580-630 nm and 650-700 nm). Pixel size was 40 nm in all dimensions and pixel dwell time was 10 μs, each line was scanned three times and the collected photon intensity signal integrated.

To calculate the effect of NR4A and NR12 photobleaching in STED imaging, a 10×10 μm^2^ area of the plasma membrane of Ptk2 cells was imaged using a 250 mW 775 nm laser power. Images were acquired every 10 s for 15 minutes. The intensity in a 2×2 μm^2^ was quantified, the background signal (as measured by the intensity in a 2×2 μm^2^ area devoid of cell signal) subtracted, and the result normalised to the intensity of the first frame. For each condition three cells were measured in each independent experiment.

Internalisation was quantified in confocal mode by acquiring 1 frame/minute for 1 h. The mean pixel intensity in cell areas that could unequivocally be identified as plasma membrane, cytosol or nucleus was measured using Fiji^58^. Intensities were normalised to that measured for the plasma membrane. GPMV formation was induced by DTT/PFA treatment of CHO cells^54^. Cells were grown on glass coverslips to a confluence of ca 75%. Cells were washed twice with GPMV buffer (2 mM CaCl2, 150 mM NaCl, 10 mM HEPES, pH 7.4), and labelled and imaged in GPMV buffer. During image acquisition, 20 mM DTT and 0.7% PFA were added. Model membrane fusion was induced by adding 10 mM CaCl_2_ to the SLB and GUV/LUV-containing chamber in HBS during image acquisition.

### FCS and STED-FCS

FCS data was acquired at an Abberior STEDYCON STED microscope mounted on a Zeiss Axio Observer Z1 inverted body, equipped with an α Plan-APOCHROMAT 100×/1.46 oil immersion objective lens (Zeiss). Samples were excited using a 40 MHz pulsed 561 nm laser, and emission stimulated by a pulsed 775 nm laser, with a repetition rate of 40 MHz. Emitted photons were collected by the objective lens, de-scanned and recorded by avalanche photodiodes with 580-625 nm and 650-700 nm filters, corresponding to the ordered and disordered channels, respectively. The signal from the APDs was cloned and sent to a Flex02-08D/C correlator card (Correlator.com). For each STED laser power, SLBs were first focused at the plane showing the highest intensity and then fine-focused (100-400 nm) to obtain maximum amplitude in FCS recordings. The focus position was adjusted after each measurement to fulfil these criteria, as small drifts of the sample would induce considerable amplitude changes in FCS curves, specially at high STED powers. FCS and STED-FCS data were acquired for 10-20 s, and for each spot and laser power at least five measurements were performed in each independent experiment. The obtained FCS curves were fitted using the FoCuS-point software^60^ to a 2D Brownian diffusion model:

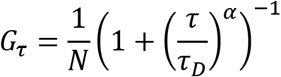

where *N* is the average number of fluorescent particles in the focal volume, *τ*_*D*_ is the average transient time and *α* the anomaly parameter. In all cases, α was close to 1, except for STED-FCS recordings at STED laser powers >100 mW, in which α = [0.80-1.00]. To quantify the resolution at different STED powers, DOPC SLBs were measured and the apparent spot size was calculated using the following equation^51^:

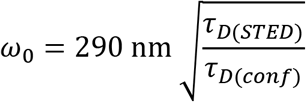

where *ω*_*0*_ is the 1/*e*^2^ radius of the gaussian beam, 290 nm is *ω*_*0*_ in confocal mode, and *τ*_*D(STED)*_ and *τ*_*D(conf)*_ are the transit times at a given STED power and in confocal mode, respectively. Counts per molecule were calculated from FCS experiments by dividing the average count rate of the measurement by the average number of molecules in the observation spot (*N*). Diffusion coefficients (*D*) were calculated from *τ*_*D*_ and *ω*_*0*_:

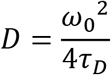

## ASSOCIATED CONTENT

### Supporting Information

The following files are available free of charge: Supplementary Figures (PDF) and Supplementary Movies 1-8 (AVI).

## AUTHOR INFORMATION

### Author Contributions

PC and ES experimental design. PC, AD and GZ data analysis. PC, AD, GZ and ES experiments. DID and ASK dye synthesis. ES and CE supervision. CE funding acquisition. The manuscript was written by PC with contributions from all authors. All authors have given approval to the final version of the manuscript.

## ACKNOWLEDGMENT

PC received funding from the European Commission Horizon 2020 Marie Skłodowska Curie programme (H2020-MSCA-IF-2019-ST project 892232 FILM-HIV) and the Basque Government (POS_2018_1_0066 and POS_2019_2_0022). We thank the Microverse Imaging Center Jena (Aurelie Jost) for providing microscope facility support. We further acknowledge funding by the Deutsche Forschungsgemeinschaft (DFG, German Research Foundation; Germany’s Excellence Strategy – EXC 2051 – Project-ID 390713860; project number 316213987 – SFB 1278; Multi-User Equipment Grant: Two-photon STED microscopy of membrane receptors), State of Thuringia (TAB)-EFRE funding (Research Group SARS-Rapid; Advanced STED microscopy), the Medical Research Council (MRC, Grant No. MC_UU_12010/unit programs G0902418 and MC_UU_12025), the EPA Cephalosporin Fund, and the John Fell Fund.

## ABBREVIATIONS

STED: stimulated-emission depletion
FCS: fluorescence correlation spectroscopy
SRM: super-resolution microscopy
SMLM: single molecule localisation microscopy
GP: generalised polarisation
GUV: giant unilamellar vesicle
SLB: supported lipid bilayer
DOPC: 1,2-dioleoyl-sn-glycero-3-phosphocholine
GPMV: giant plasma membrane vesicle
DOPE: 1,2-dioleoyl-sn-glycero-3-phosphoethanolamine
DOPS: 1,2-dioleoyl-sn-glycero-3-phosphoserine
POPC: 1-palmitoyl-2-oleoyl-glycero-3-phosphocholine
LUV: large unilamellar vesicle

## REFERENCES

(1) Sezgin, E.; Levental, I.; Mayor, S.; Eggeling, C. The Mystery of Membrane Organization: Composition, Regulation and Roles of Lipid Rafts. Nature Reviews Molecular Cell Biology 2017, 18 (6), 361–374. https://doi.org/10.1038/nrm.2017.16.

(2) Levental, K. R.; Surma, M. A.; Skinkle, A. D.; Lorent, J. H.; Zhou, Y.; Klose, C.; Chang, J. T.; Hancock, J. F.; Levental, I. W-3 Polyunsaturated Fatty Acids Direct Differentiation of the Membrane Phenotype in Mesenchymal Stem Cells to Potentiate Osteogenesis. Science Advances 2017, 16.

(3) Favard, C.; Chojnacki, J.; Merida, P.; Yandrapalli, N.; Mak, J.; Eggeling, C.; Muriaux, D. HIV-1 Gag Specifically Restricts PI(4,5)P2 and Cholesterol Mobility in Living Cells Creating a Nanodomain Platform for Virus Assembly. Science Advances 2019, 5, eaaw8651.

(4) Sengupta, P.; Seo, A. Y.; Pasolli, H. A.; Song, Y. E.; Johnson, M.; Lippincott-Schwartz, J. A Lipid-Based Partitioning Mechanism for Selective Incorporation of Proteins into Membranes of HIV Particles. Nature Cell Biology 2019, 21 (4), 452–461. https://doi.org/10.1038/s41556-019-0300-y.

(5) Eggeling, C.; Ringemann, C.; Medda, R.; Schwarzmann, G.; Sandhoff, K.; Polyakova, S.; Belov, V. N.; Hein, B.; von Middendorff, C.; Schönle, A.; Hell, S. W. Direct Observation of the Nanoscale Dynamics of Membrane Lipids in a Living Cell. Nature 2009, 457 (7233), 1159–1162. https://doi.org/10.1038/nature07596.

(6) Schwille, P.; Korlach, J.; Webb, W. W. Fluorescence Correlation Spectroscopy with Single-Molecule Sensitivity on Cell and Model Membranes. Cytometry 1999, 36 (3), 176–182. https://doi.org/10.1002/(SICI)1097-0320(19990701)36:3176::AID-CYTO53.0.CO;2-F.

(7) Yu, W.; So, P. T.; French, T.; Gratton, E. Fluorescence Generalized Polarization of Cell Membranes: A Two-Photon Scanning Microscopy Approach. Biophysical Journal 1996, 70 (2), 626–636. https://doi.org/10.1016/S0006-3495(96)79646-7.

(8) Colom, A.; Derivery, E.; Soleimanpour, S.; Tomba, C.; Molin, M. D.; Sakai, N.; González-gaitán, M.; Matile, S.; Roux, A. A Fluorescent Membrane Tension Probe. Nature Chemistry 2018, 10, 1118–1125. https://doi.org/10.1038/s41557-018-0127-3.

(9) Sezgin, E. Super-Resolution Optical Microscopy for Studying Membrane Structure and Dynamics. Journal of Physics: Condensed Matter 2017, 29 (27), 273001. https://doi.org/10.1088/1361-648X/aa7185.

(10) Betzig, E.; Patterson, G. H.; Sougrat, R.; Lindwasser, O. W.; Olenych, S.; Bonifacino, J. S.; Davidson, M. W.; Lippincott-Schwartz, J.; Hess, H. F. Imaging Intracellular Fluorescent Proteins at Nanometer Resolution. Science 2006, 313 (5793), 1642–1645. https://doi.org/10.1126/science.1127344.

(11) Balzarotti, F.; Eilers, Y.; Gwosch, K. C.; Gynnå, A. H.; Westphal, V.; Stefani, F. D.; Elf, J.; Hell, S. W. Nanometer Resolution Imaging and Tracking of Fluorescent Molecules with Minimal Photon Fluxes. Science 2017, 355 (February), 606–612. https://doi.org/10.1126/science.aak9913.

(12) Vicidomini, G.; Bianchini, P.; Diaspro, A. STED Super-Resolved Microscopy. Nature Methods 2018, 15, 173–182. https://doi.org/10.1038/nmeth.4593.

(13) Hedde, P. N.; Dörlich, R. M.; Blomley, R.; Gradl, D.; Oppong, E.; Cato, A. C. B.; Nienhaus, G. U. Stimulated Emission Depletion-Based Raster Image Correlation Spectroscopy Reveals Biomolecular Dynamics in Live Cells. Nature Communications 2013, 4 (1), 2093. https://doi.org/10.1038/ncomms3093.

(14) Sezgin, E.; Schneider, F.; Zilles, V.; Urbančič, I.; Garcia, E.; Waithe, D.; Klymchenko, A. S.; Eggeling, C. Polarity-Sensitive Probes for Superresolution Stimulated Emission Depletion Microscopy. Biophysical Journal 2017, 113 (6), 1321–1330. https://doi.org/10.1016/j.bpj.2017.06.050.

(15) Barbotin, A.; Urbančič, I.; Galiani, S.; Eggeling, C.; Booth, M.; Sezgin, E. Z-STED Imaging and Spectroscopy to Investigate Nanoscale Membrane Structure and Dynamics. Biophysical Journal 2020, 118 (10), 2448–2457. https://doi.org/10.1016/j.bpj.2020.04.006.

(16) Eggeling, C.; Willig, K. I.; Sahl, S. J.; Hell, S. W. Lens-Based Fluorescence Nanoscopy. Quart. Rev. Biophys. 2015, 48 (2), 178–243. https://doi.org/10.1017/S0033583514000146.

(17) Donnert, G.; Keller, J.; Medda, R.; Andrei, M. A.; Rizzoli, S. O.; Lührmann, R.; Jahn, R.; Eggeling, C.; Hell, S. W. Macromolecular-Scale Resolution in Biological Fluorescence Microscopy. PNAS 2006, 103 (31), 11440–11445. https://doi.org/10.1073/pnas.0604965103.

(18) Oracz, J.; Westphal, V.; Radzewicz, C.; Sahl, S. J.; Hell, S. W. Photobleaching in STED Nanoscopy and Its Dependence on the Photon Flux Applied for Reversible Silencing of the Fluorophore. Sci Rep 2017, 7 (1), 11354. https://doi.org/10.1038/s41598-017-09902-x.

(19) Song, L.; Varma, C. A.; Verhoeven, J. W.; Tanke, H. J. Influence of the Triplet Excited State on the Photobleaching Kinetics of Fluorescein in Microscopy. Biophys J 1996, 70 (6), 2959–2968. https://doi.org/10.1016/S0006-3495(96)79866-1.

(20) Eggeling, C.; Windergren, J.; Rigler, R.; Seidel, C. A. M. Photobleaching of Fluorescent Dyes under Conditions Used for Single-Molecule Detection: Evidence of Two-Step Photolysis. Analytical Chemistry 1998, 70 (13), 2651–2659. https://doi.org/10.1021/ac980027p.

(21) Hotta, J.; Fron, E.; Dedecker, P.; Janssen, K. P. F.; Li, C.; Müllen, K.; Harke, B.; Bückers, J.; Hell, S. W.; Hofkens, J. Spectroscopic Rationale for Efficient Stimulated-Emission Depletion Microscopy Fluorophores. J. Am. Chem. Soc. 2010, 132 (14), 5021–5023. https://doi.org/10.1021/ja100079w.

(22) Kolmakov, K.; Belov, V. N.; Bierwagen, J.; Ringemann, C.; Müller, V.; Eggeling, C.; Hell, S. W. Red-Emitting Rhodamine Dyes for Fluorescence Microscopy and Nanoscopy. Chemistry 2010, 16 (1), 158–166. https://doi.org/10.1002/chem.200902309.

(23) Zheng, Q.; Lavis, L. D. Development of Photostable Fluorophores for Molecular Imaging. Current Opinion in Chemical Biology 2017, 39, 32–38. https://doi.org/10.1016/j.cbpa.2017.04.017.

(24) Donnert, G.; Eggeling, C.; Hell, S. W. Major Signal Increase in Fluorescence Microscopy through Dark-State Relaxation. Nature Methods 2007, 4 (1), 81–86. https://doi.org/10.1038/nmeth986.

(25) Wu, Y.; Wu, X.; Lu, R.; Zhang, J.; Toro, L.; Stefani, E. Resonant Scanning with Large Field of View Reduces Photobleaching and Enhances Fluorescence Yield in STED Microscopy. Scientific Reports 2015, 5 (1), 14766. https://doi.org/10.1038/srep14766.

(26) Donnert, G.; Eggeling, C.; Hell, S. W. Triplet-Relaxation Microscopy with Bunched Pulsed Excitation. Photochemical and Photobiological Sciences 2009, 8, 481–485. https://doi.org/10.1039/b903357m.

(27) Heine, J.; Reuss, M.; Harke, B.; D’Este, E.; Sahl, S. J.; Hell, S. W. Adaptive-Illumination STED Nanoscopy. PNAS 2017, 114 (37), 9797–9802. https://doi.org/10.1073/pnas.1708304114.

(28) Sharonov, A.; Hochstrasser, R. M. Wide-Field Subdiffraction Imaging by Accumulated Binding of Diffusing Probes. Proceedings of the National Academy of Sciences of the United States of America 2006, 103 (50), 18911–18916. https://doi.org/10.1073/pnas.0609643104.

(29) Jungmann, R.; Avendaño, M. S.; Woehrstein, J. B.; Dai, M.; Shih, W. M.; Yin, P. Multiplexed 3D Cellular Super-Resolution Imaging with DNA-PAINT and Exchange-PAINT. Nature Methods 2014, 11 (3), 313–318. https://doi.org/10.1038/nmeth.2835.

(30) Feng Gao; Erwen Mei; Manho Lim, † and; Hochstrasser*, R. M. Probing Lipid Vesicles by Bimolecular Association and Dissociation Trajectories of Single Molecules https://pubs.acs.org/doi/pdf/10.1021/ja058098a (accessed Mar 19, 2021). https://doi.org/10.1021/ja058098a.

(31) Bongiovanni, M. N.; Godet, J.; Horrocks, M. H.; Tosatto, L.; Carr, A. R.; Wirthensohn, D. C.; Ranasinghe, R. T.; Lee, J. E.; Ponjavic, A.; Fritz, J. V.; Dobson, C. M.; Klenerman, D.; Lee, S. F. Multi-Dimensional Super-Resolution Imaging Enables Surface Hydrophobicity Mapping. Nature Communications 2016, 7, 1–9. https://doi.org/10.1038/ncomms13544.

(32) Moon, S.; Yan, R.; Kenny, S. J.; Shyu, Y.; Xiang, L.; Li, W.; Xu, K. Spectrally Resolved, Functional Super-Resolution Microscopy Reveals Nanoscale Compositional Heterogeneity in Live-Cell Membranes. Journal of the American Chemical Society 2017, 139 (32), 10944–10947. https://doi.org/10.1021/jacs.7b03846.

(33) Yan, R.; Chen, K.; Xu, K. Probing Nanoscale Diffusional Heterogeneities in Cellular Membranes through Multidimensional Single-Molecule and Super-Resolution Microscopy. J. Am. Chem. Soc. 2020, 142 (44), 18866–18873. https://doi.org/10.1021/jacs.0c08426.

(34) Danylchuk, D. I.; Moon, S.; Xu, K.; Klymchenko, A. S. Tailor-made Switchable Solvatochromic Probes for Live-cell Super-resolution Imaging of Plasma Membrane Organization. Angewandte Chemie 2019, 131, 15062–15066. https://doi.org/10.1002/ange.201907690.

(35) Spahn, C.; Grimm, J. B.; Lavis, L. D.; Lampe, M.; Heilemann, M. Whole-Cell, 3D, and Multicolor STED Imaging with Exchangeable Fluorophores. Nano Letters 2019, 19 (1), 500–505. https://doi.org/10.1021/acs.nanolett.8b04385.

(36) Sackett, D. L.; Knutson, J. R.; Wolff, J. Hydrophobic Surfaces of Tubulin Probed by Time-Resolved and Steady-State Fluorescence of Nile Red. Journal of Biological Chemistry 1990, 265 (25), 14899–14906. https://doi.org/10.1016/S0021-9258(18)77201-3.

(37) Hou, Y.; Bardo, A. M.; Martinez, C.; Higgins, D. A. Characterization of Molecular Scale Environments in Polymer Films by Single Molecule Spectroscopy. J. Phys. Chem. B 2000, 104 (2), 212–219. https://doi.org/10.1021/jp992312y.

(38) Reichardt, C. Solvatochromic Dyes as Solvent Polarity Indicators. Chem. Rev. 1994, 94 (8), 2319–2358. https://doi.org/10.1021/cr00032a005.

(39) Klymchenko, A. S. Solvatochromic and Fluorogenic Dyes as Environment-Sensitive Probes: Design and Biological Applications. Acc. Chem. Res. 2017, 50 (2), 366–375. https://doi.org/10.1021/acs.accounts.6b00517.

(40) Parasassi, T.; De Stasio, G.; d’Ubaldo, A.; Gratton, E. Phase Fluctuation in Phospholipid Membranes Revealed by Laurdan Fluorescence. Biophysical Journal 1990, 57 (6), 1179–1186. https://doi.org/10.1016/S0006-3495(90)82637-0.

(41) Klymchenko, A. S.; Kreder, R. Fluorescent Probes for Lipid Rafts: From Model Membranes to Living Cells. Chemistry & Biology 2014, 21 (1), 97–113. https://doi.org/10.1016/j.chembiol.2013.11.009.

(42) Carravilla, P.; Nieva, J. L.; Goñi, F. M.; Requejo-Isidro, J.; Huarte, N. Two-Photon Laurdan Studies of the Ternary Lipid Mixture DOPC:SM:Cholesterol Reveal a Single Liquid Phase at Sphingomyelin:Cholesterol Ratios Lower Than 1. Langmuir 2015, 31 (9), 2808–2817. https://doi.org/10.1021/la504251u.

(43) Jin, L.; Millard, A. C.; Wuskell, J. P.; Clark, H. A.; Loew, L. M. Cholesterol-Enriched Lipid Domains Can Be Visualized by Di-4-ANEPPDHQ with Linear and Nonlinear Optics. Biophysical Journal 2005, 89 (1), L04–L06. https://doi.org/10.1529/biophysj.105.064816.

(44) Jin, L.; Millard, A. C.; Wuskell, J. P.; Dong, X.; Wu, D.; Clark, H. A.; Loew, L. M. Characterization and Application of a New Optical Probe for Membrane Lipid Domains. Biophysical Journal 2006, 90 (7), 2563–2575. https://doi.org/10.1529/biophysj.105.072884.

(45) Danylchuk, D. I.; Sezgin, E.; Chabert, P.; Klymchenko, A. S. Redesigning Solvatochromic Probe Laurdan for Imaging Lipid Order Selectively in Cell Plasma Membranes. Analytical Chemistry 2020, 92, 14798–14805. https://doi.org/10.1021/acs.analchem.0c03559.

(46) Kucherak, O. A.; Oncul, S.; Darwich, Z.; Yushchenko, D. A.; Arntz, Y.; Didier, P.; Mély, Y.; Klymchenko, A. S. Switchable Nile Red-Based Probe for Cholesterol and Lipid Order at the Outer Leaflet of Biomembranes. Journal of the American Chemical Society 2010, 132 (13), 4907–4916. https://doi.org/10.1021/ja100351w.

(47) Owen, D. M.; Rentero, C.; Magenau, A.; Abu-Siniyeh, A.; Gaus, K. Quantitative Imaging of Membrane Lipid Order in Cells and Organisms. Nature Protocols 2011, 7 (1), 24–35. https://doi.org/10.1038/nprot.2011.419.

(48) Sezgin, E.; Waithe, D.; Bernardino De La Serna, J.; Eggeling, C. Spectral Imaging to Measure Heterogeneity in Membrane Lipid Packing. ChemPhysChem 2015, 16 (7), 1387–1394. https://doi.org/10.1002/cphc.201402794.

(49) Eggeling, C.; Volkmer, A.; Seidel, C. A. M. Molecular Photobleaching Kinetics of Rhodamine 6G by One- and Two-Photon Induced Confocal Fluorescence Microscopy. ChemPhysChem 2005, 6 (5), 791–804. https://doi.org/10.1002/cphc.200400509.

(50) van Meer, G.; Voelker, D. R.; Feigenson, G. W. Membrane Lipids: Where They Are and How They Behave. Nature Reviews Molecular Cell Biology 2008, 9 (2), 112–124. https://doi.org/10.1038/nrm2330.

(51) Sezgin, E.; Schneider, F.; Galiani, S.; Urbančič, I.; Waithe, D.; Lagerholm, B. C.; Eggeling, C. Measuring Nanoscale Diffusion Dynamics in Cellular Membranes with Super-Resolution STED–FCS. Nature Protocols 2019, 14, 1054–1083. https://doi.org/10.1038/s41596-019-0127-9.

(52) Urbančič, I.; Brun, J.; Shrestha, D.; Waithe, D.; Eggeling, C.; Chojnacki, J. Lipid Composition but Not Curvature Is the Determinant Factor for the Low Molecular Mobility Observed on the Membrane of Virus-like Vesicles. Viruses 2018, 10 (8). https://doi.org/10.3390/v10080415.

(53) Klar, T. A.; Jakobs, S.; Dyba, M.; Egner, A.; Hell, S. W. Fluorescence Microscopy with Diffraction Resolution Barrier Broken by Stimulated Emission. PNAS 2000, 97 (15), 8206–8210. https://doi.org/10.1073/pnas.97.15.8206.

(54) Sezgin, E.; Kaiser, H.-J.; Baumgart, T.; Schwille, P.; Simons, K.; Levental, I. Elucidating Membrane Structure and Protein Behavior Using Giant Plasma Membrane Vesicles. Nature Protocols 2012, 7 (6), 1042–1051. https://doi.org/10.1038/nprot.2012.059.

(55) Skinkle, A. D.; Levental, K. R.; Levental, I. Cell-Derived Plasma Membrane Vesicles Are Permeable to Hydrophilic Macromolecules. Biophys J 2020, 118 (6), 1292–1300. https://doi.org/10.1016/j.bpj.2019.12.040.

(56) Wilschut, J.; Duezguenes, N.; Papahadjopoulos, D. Calcium/Magnesium Specificity in Membrane Fusion: Kinetics of Aggregation and Fusion of Phosphatidylserine Vesicles and the Role of Bilayer Curvature. Biochemistry 1981, 20 (11), 3126–3133. https://doi.org/10.1021/bi00514a022.

(57) Kilian, N.; Goryaynov, A.; Lessard, M. D.; Hooker, G.; Toomre, D.; Rothman, J. E.; Bewersdorf, J. Assessing Photodamage in Live-Cell STED Microscopy. Nature Methods 2018, 15 (10), 755–756. https://doi.org/10.1038/s41592-018-0145-5.

(58) Schindelin, J.; Arganda-Carreras, I.; Frise, E.; Kaynig, V.; Longair, M.; Pietzsch, T.; Preibisch, S.; Rueden, C.; Saalfeld, S.; Schmid, B.; Tinevez, J.-Y.; White, D. J.; Hartenstein, V.; Eliceiri, K.; Tomancak, P.; Cardona, A. Fiji: An Open-Source Platform for Biological-Image Analysis. Nature Methods 2012, 9 (7), 676–682. https://doi.org/10.1038/nmeth.2019.

(59) Lord, S. J.; Velle, K. B.; Mullins, R. D.; Fritz-Laylin, L. K. SuperPlots: Communicating Reproducibility and Variability in Cell Biology. Journal of Cell Biology 2020, 219 (e202001064). https://doi.org/10.1083/jcb.202001064.

(60) Waithe, D.; Clausen, M. P.; Sezgin, E.; Eggeling, C. FoCuS-Point: Software for STED Fluorescence Correlation and Time-Gated Single Photon Counting. Bioinformatics 2016, 32 (6), 958–960. https://doi.org/10.1093/bioinformatics/btv687.

