## Supplementary Figures for "STED super-resolution imaging of membrane packing and dynamics by exchangeable polarity-sensitive dyes"

### Real-time quantification of membrane packing and dynamics using exchangeable polarity-sensitive dyes and STED super-resolution microscopy

*Pablo Carravilla[1,2*], Anindita Dasgupta[1,2], Gaukhar Zhurgenbayeva[2,3], Dmytro I. Danylchuk[4], Andrey S. Klymchenko[4], Erdinc Sezgin[5]* and Christian Eggeling[1,2,3,6,7]*

[1]Leibniz Institute of Photonic Technology, 07745 Jena, Germany

[2]Institute of Applied Optics and Biophysics, Friedrich-Schiller-University Jena, 07743 Jena, Germany

[3]Jena School for Microbial Communication (JSMC), Friedrich-Schiller-University Jena, 07743 Jena, Germany

[4]Laboratoire de Bioimagerie et Pathologies, UMR 7021 CNRS, Université de Strasbourg, 67401 Illkirch, France

[5]Science for Life Laboratory, Department of Women's and Children's Health, Karolinska Institutet, 17177 Stockholm, Sweden

[6]MRC Human Immunology Unit, Weatherall Institute of Molecular Medicine, University of Oxford, OX39DS Oxford, UK

[7]Jena Center for Soft Matter (JCSM), 07743 Jena, Germany

*To whom correspondence should be addressed

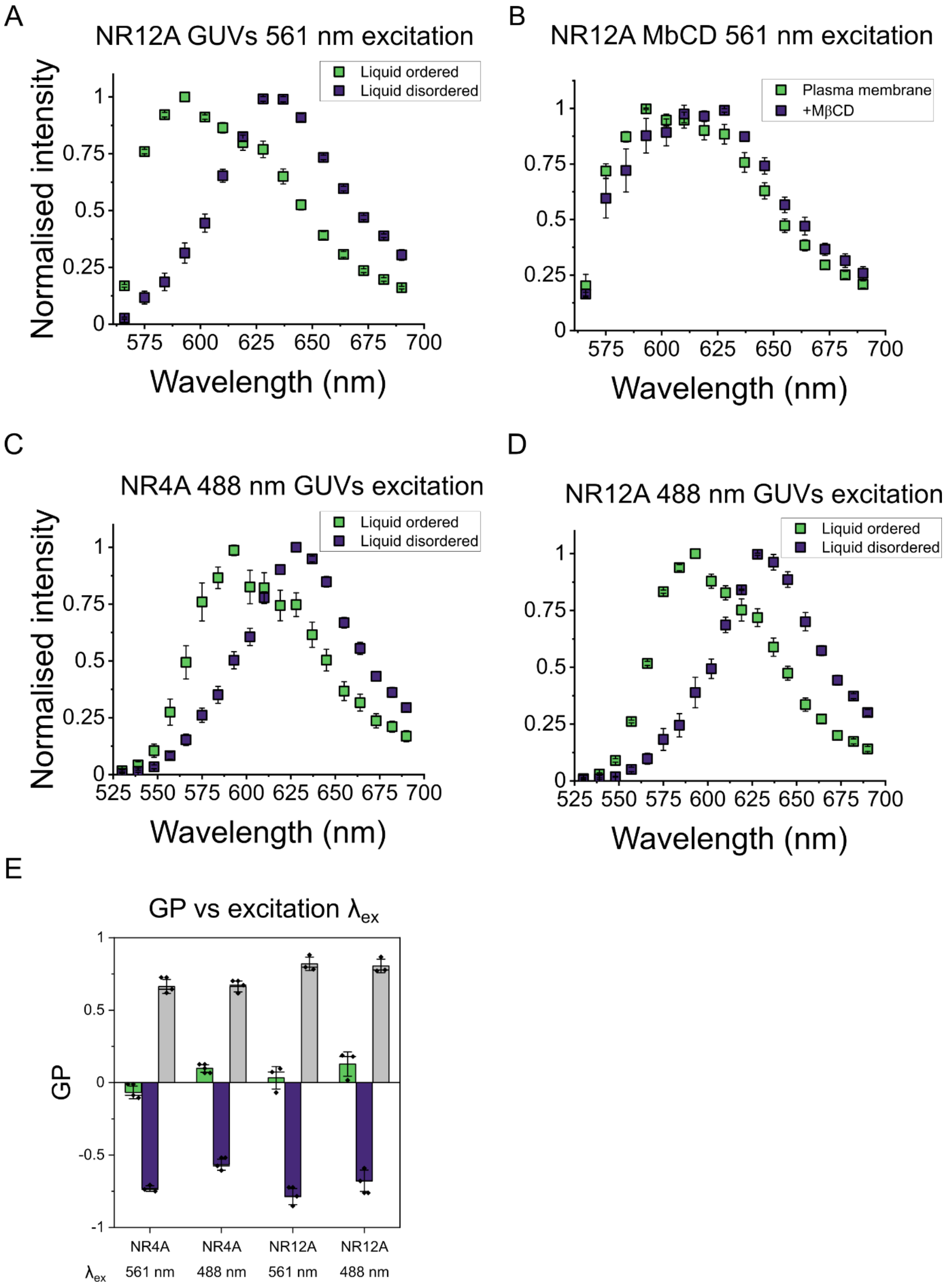


**Supplementary Figure 1. Characterisation of NR4A and NR12A spectral properties.** (A) Emission spectra of NR12A in phase separated GUVs. Conditions as in Figure 1C. (B) Emission spectra of NR12A in control and MβCD-treated C2BBe1 cells. Conditions as in Figure 1E. (C) and (D) emission spectra of NR4A and NR12A, respectively in phase-separated GUVs under 488 nm excitation. 561 nm excitation was preferred since it yielded a higher intensity signal (data not shown). Conditions as in Figure 1C. (E) GP-resolution comparison between 488 nm and 561 nm excitation of NR4A and NR12A in phase-separated GUVs. Conditions as in Figure 1D.

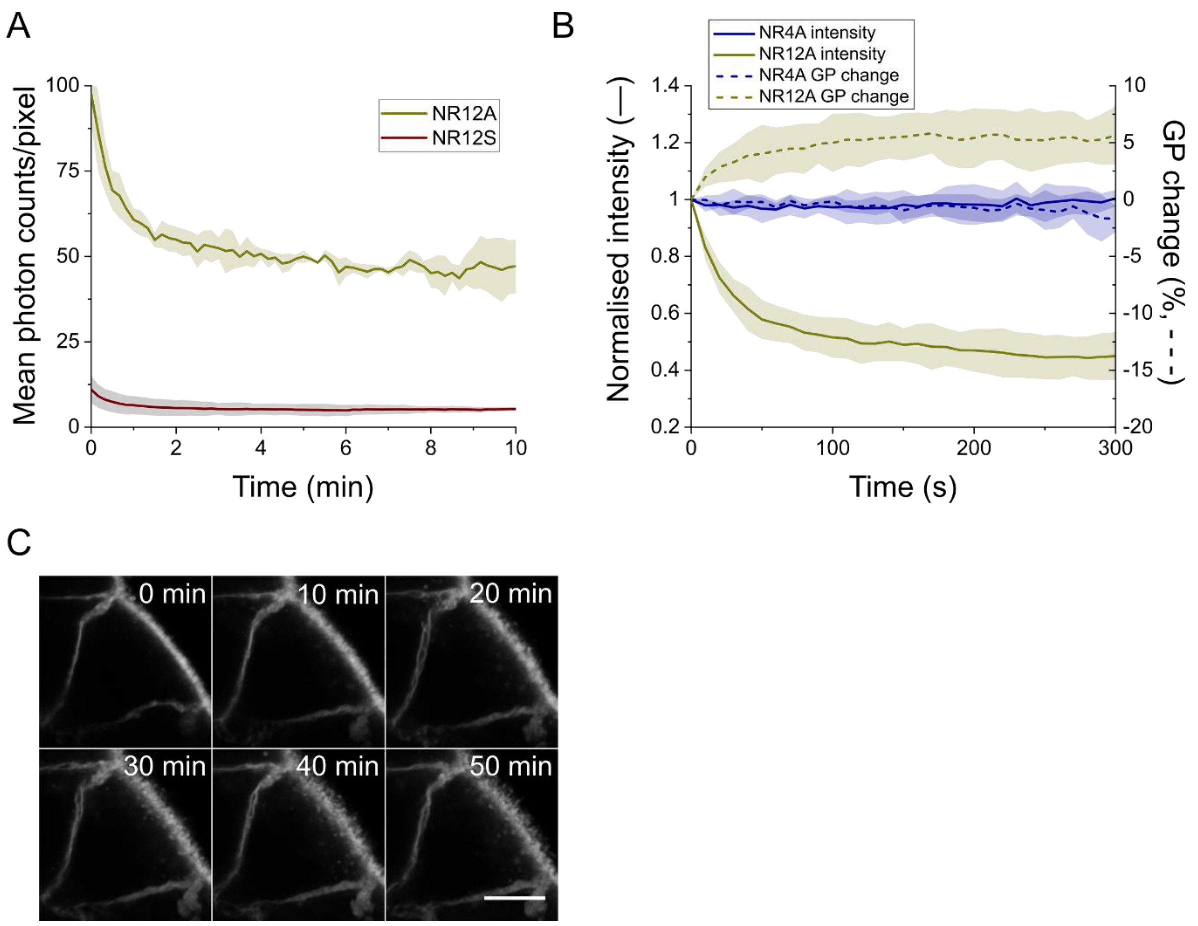


**Supplementary Figure 2. Plasma membrane STED imaging of polarity-sensitive dyes NR4A, NR12A and NR12S.** (A) Raw intensity signal comparison of Ptk2 plasma membrane signal between NR12S, a previous generation Nile Red derivative, and NR12A. (B) STED laser-induced photobleaching causes a ca. 5% change in the measured GP value in Ptk2 plasma membranes. (C) Micrographs of confocal images of C2BBe1 cells labelled with NR12A to measure dye internalisation. Internalisation quantification is shown in Figure 2D. Otherwise, conditions as in Figure 2.

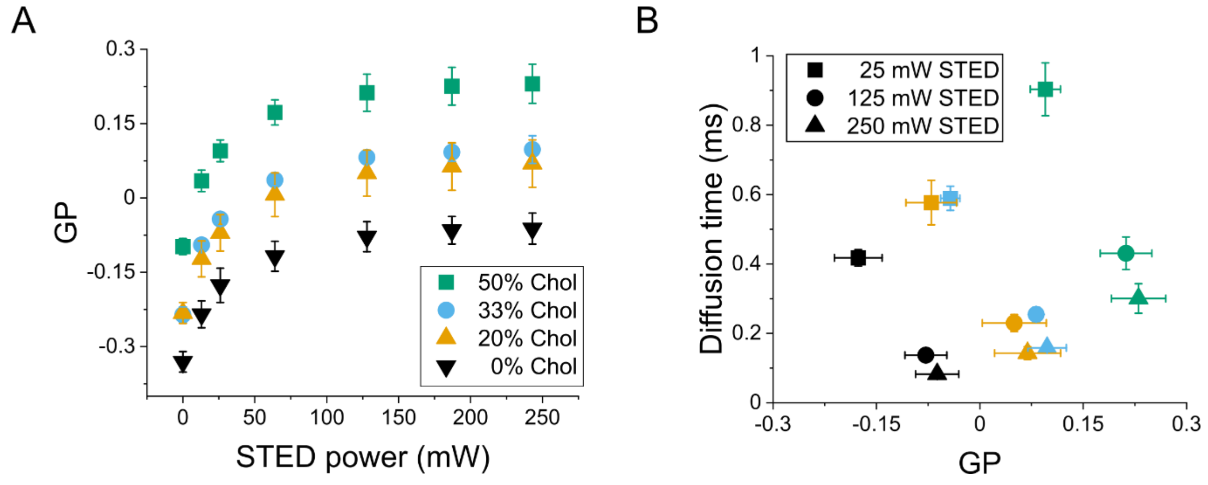


**Supplementary Figure 3. STED-FCS measurements of NR4A.** (A) NR4A GP values measured in cholesterol-containing SLBs. GP values depend on the STED laser power, but show a high sensitivity at all measured powers. (B) Dynamics and lipid packing correlate independently of the STED laser power used. Colour codes as in panel A. Conditions as in Figure 3D.

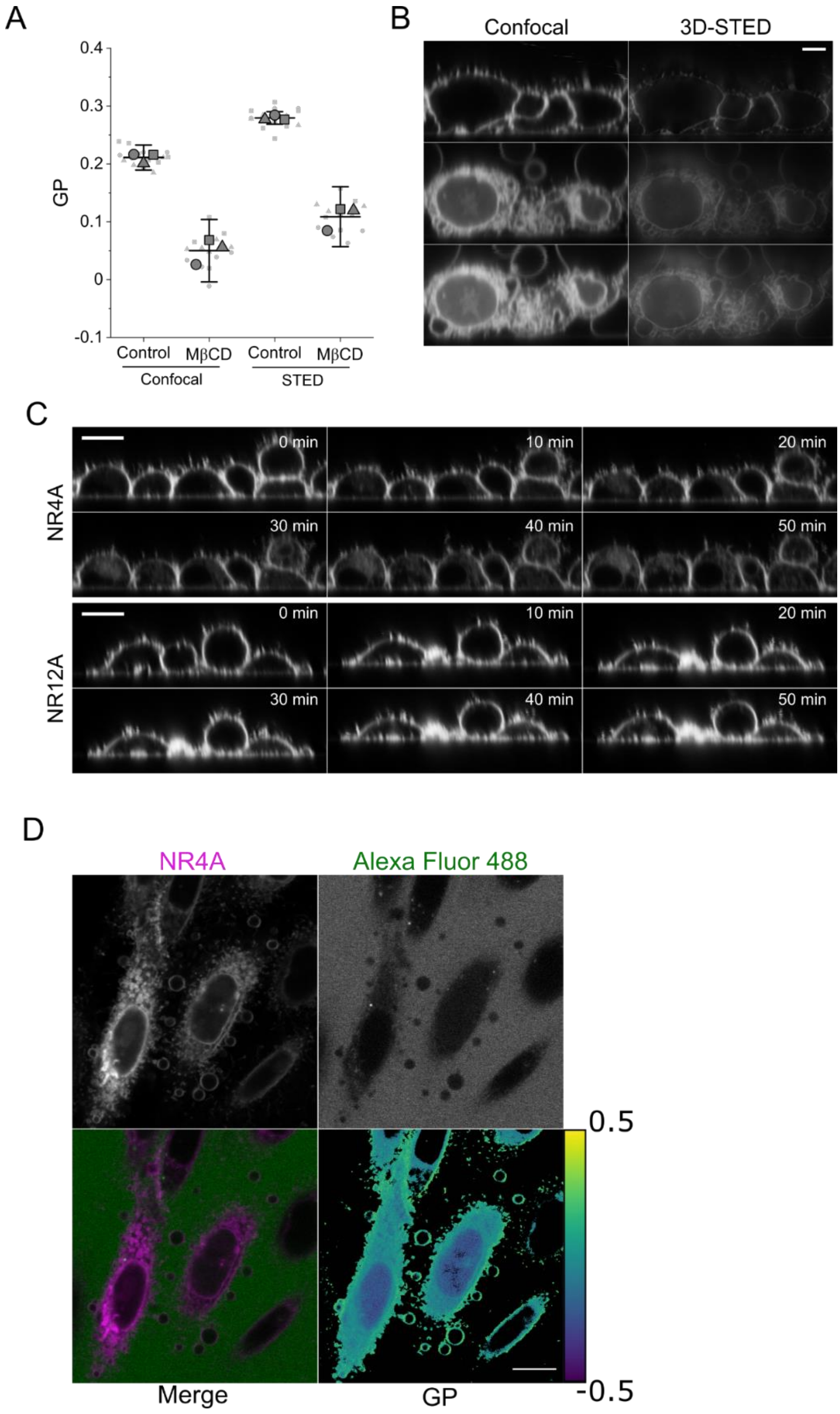


**Supplementary Figure 4. NR4A imaging of live cell plasma membranes.** (A) Comparison between confocal and STED-GP values of control and MβCD-treated (20 mM, 1h, 37 °C) C2BBe1 cells. STED laser power was 250 mW. Conditions as in Figure 4A. (B) Plasma membrane vesiculation of 293T/17 cells induced by DTT/PFA treatment at 37 °C. Conditions as in Figure 4C. Scale bar is 5 µm. (C) Micrographs of confocal *xz* images of the internalisation of NR4A (top) and NR12A (bottom) in CHO cells. Scale bar is 10 µm. (D) Micrographs of CHO cells after 30-minute DTT/PFA treatment. DTT/PFA induce internalisation of NR4A (magenta, 500 nM), but not of the small hydrophilic Alexa Fluor 488 (green, 1 µM). GP images shows the lower packing of internal cell membranes.

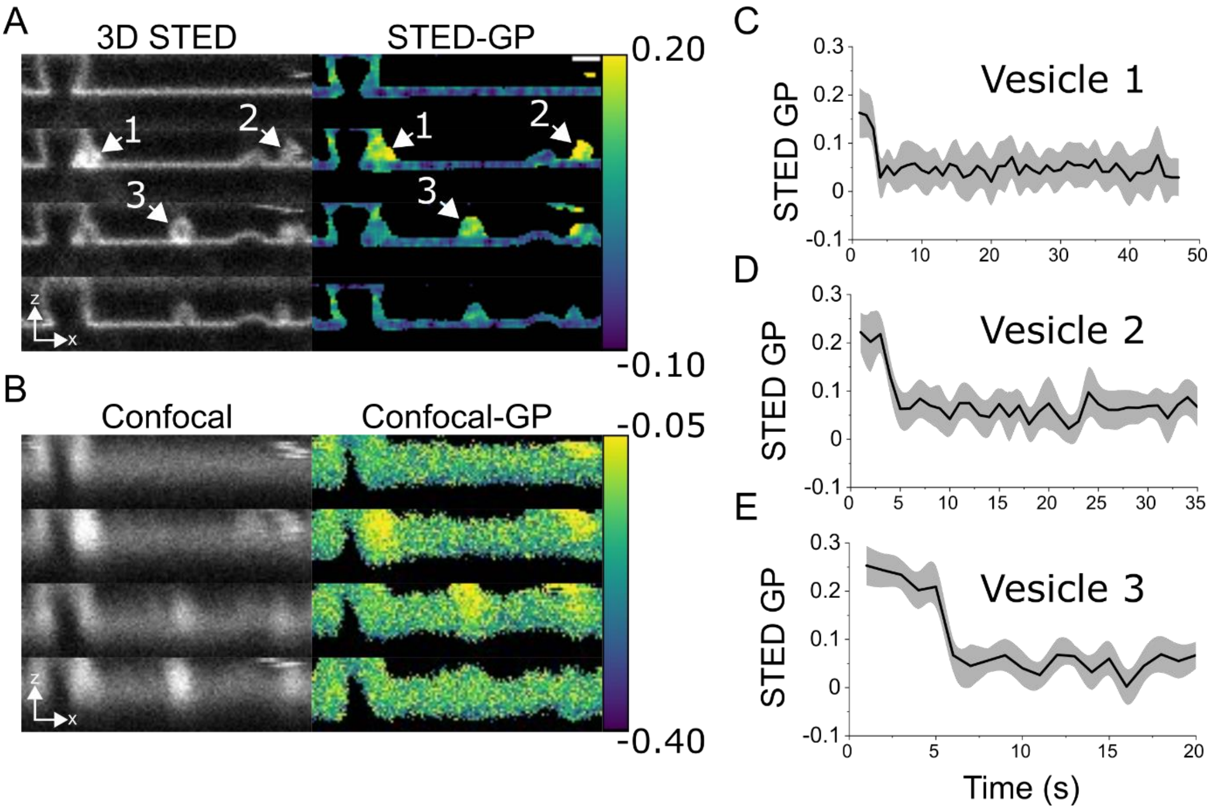


**Supplementary Figure 5. Real time visualisation of LUV fusion with an SLB.** (A) 3D-STED intensity (left) and GP (right) images of POPC:cholesterol (2:1 mol ratio) LUVs fusing with a DOPC:DOPE:DOPS (4:3:3 mol ratio) bilayer. Scale bar is 500 nm. (B) Same as in A, but acquired in confocal mode. (C-E) Quantification of the GP changes over time of the vesicles marked with an arrowhead in panel A. Otherwise conditions as in Figure 5.
